# Sequestration of histidine kinases by non-cognate response regulators establishes a threshold level of stimulation for bacterial two-component signaling

**DOI:** 10.1101/2021.12.30.474508

**Authors:** Gaurav D. Sankhe, Rubesh Raja, Narendra M. Dixit, Deepak Kumar Saini

## Abstract

Two-component signaling systems (TCSs) in bacteria are often positively auto-regulated, where the histidine kinase (HK) and response regulator (RR) proteins comprising a TCS are expressed downstream of the signal they transduce. This auto-regulation improves the sensitivity of the TCS to stimuli and amplifies adaptive responses. The downside, however, is that the TCS may mount disproportionately large responses to weak or fleeting signals. How bacteria prevent such disproportionate responses is not known. Here, we show that sequestration of phosphorylated HKs by non-cognate RRs serves as a design to prevent such disproportionate responses. Using TCSs of *M. tuberculosis* as model systems, we found that with every one of the five HKs we studied, there was at least one non-cognate RR with higher affinity than that of the cognate RR for the HK. Phosphorylated HKs would thus preferentially bind the non-cognate RRs, suppressing signal transduction through the cognate pathways, which we demonstrated *in vitro*. Using mathematical modeling of TCS signaling *in vivo*, we predicted that this sequestration would introduce a threshold level of stimulation for a significant response, preventing responses to signals below this threshold. Finally, we showed *in vivo* using tunable expression systems in *M. bovis* that upregulation of a higher affinity non-cognate RR substantially suppressed the output from the cognate TCS pathway, presenting strong evidence of sequestration by non-cognate RRs as a design to regulate TCS signaling. Blocking this sequestration may be a novel intervention strategy, as it would compromise bacterial fitness by letting it respond unnecessarily to signals.

## INTRODUCTION

Two-component signaling systems (TCSs) form the primary apparati in bacteria for sensing and responding to extracellular cues.^1^ Bacteria can have a few tens to a few hundred distinct TCSs. Each TCS comprises a sensor histidine kinase (HK), which is usually a transmembrane protein with a variable sensory domain and conserved catalytic domains, and a cognate cytosolic response regulator (RR) protein which also contains a conserved catalytic domain and a variable output domain.^1,2^ Stimulation by an extracellular cue through the input domain leads to HK autophosphorylation, followed by phosphotransfer from the HK to its cognate RR, which involves interaction between the conserved catalytic domains of both the proteins. The phosphorylation alters the DNA binding properties of the variable output domain of the RR, resulting in transcriptional changes of downstream genes and an adaptive response to the external stimulus.^1,2^

An important feature of the adaptive response, prevalent across TCSs, is positive autoregulation^3^: The phosphorylated RR upregulates the expression of the corresponding HK and RR proteins. The increased HK and RR levels can increase the sensitivity of the TCS to the external stimulus and the magnitude of the adaptive response, respectively.^4^ When the external stimulus is strong and persistent, positive autoregulation confers an advantage on the bacteria by expediting and amplifying the adaptive response. Indeed, positive autoregulation is a widely recognized biological design for amplifying responses, in addition to its effects on promoting step-like responses, hysteresis, and memory.^5-8^ When the stimulus is weak or fleeting, however, positive autoregulation can be a disadvantage as it can lead to the mounting of a response that is disproportionately amplified and sustained given the stimulus. An important question that follows is how bacteria guard themselves against such disproportionate responses despite the presence of positive autoregulation.

Recent studies have presented designs that could create the requirement of a threshold level of stimulation before a response is mounted. For instance, in some plant TCSs and, more recently, in the TCS ArcAB of *E. coli*, multiple intermediate phosphotransfer events, resulting in a phosphorelay, have been identified in the phosphotransfer from HK to the cognate RR.^9,10^ During the process, it is conceivable that a decay of a short-lived stimulus would cause the deactivation of HK and abort phosphotransfer. A minimum strength and/or duration of stimulation is thus necessary to complete the signal transduction event. Similarly, negative feedback could prevent the autoregulatory loop from getting activated until a threshold level of stimulation is realized.^11,12^ Other designs include scaffolds that bind HK and dampen its activity^13,14^, small molecules that allosterically increase the phosphatase activity of HKs leading to rapid dephosphorylation of RRs following phosphotransfer^15-17^, and phosphate sinks that prematurely terminate signaling^18^. Indeed, some of these designs are being explored as novel routes to actively tune the detection thresholds of TCSs.^16^ The designs explain the emergence of thresholds but imply that different designs exist in different settings for achieving the same goal. Given the ubiquitous nature of TCSs, we reasoned that a more widely prevalent motif for preventing disproportionate responses, both in strength and duration, may exist in bacterial systems.

Here, we identify a new design principle, involving the sequestration of autophosphorylated HKs by non-cognate RRs, that precludes an amplified response unless stimulation crosses a threshold level. We reasoned that if a phosphorylated HK has a higher binding affinity for a non-cognate RR than its cognate counterpart, then the HK is likely to be sequestered by the non-cognate RR. Only if the external stimulus leads to the accumulation of a sufficient amount of the phosphorylated HKs will the sequestration be overcome and phosphotransfer to the cognate RR occur. We unravel this design using mycobacterial TCSs, which we studied under *in vitro* and *in vivo* conditions, and using mathematical modeling. We found that the design exists in every one of the TCSs we studied, suggesting that it may be a widely prevalent mechanism for fine-tuning TCS signaling.

## RESULTS

Sequestration would occur if phosphorylated HKs were to bind preferentially to non-cognate RRs, so that their ability to transfer phosphoryl groups to their cognate RR partners is compromised. TCSs of *M. tuberculosis* have been shown *in vitro* to exhibit extensive cross-talk, where phosphorylated HKs not only bind but also transfer phosphoryl groups to non-cognate RRs.^19,20^ They thus offered an excellent model system for us to test our hypothesis.

### The phosphorylated HK MtrB has a higher affinity for some non-cognate RRs than its cognate RR MtrA

We considered first the TCS MtrAB, which is among the most promiscuous of the TCSs of *M. tuberculosis*^19^ and is known to exhibit positive autoregulation^21^. We measured the binding affinities of the phosphorylated, GFP-tagged HK MtrB, reported to be active in previous studies^22^, for its cognate RR MtrA and several non-cognate RRs, using microscale thermophoresis (Methods). We found that the equilibrium dissociation constant, *K*_*D*_, of the phosphorylated MtrB for MtrA was 268±14 nM, whereas it was 80±4 nM and 98±5 nM, for the non-cognate RRs NarL and TcrX, respectively (Figure 1). Remarkably, the phosphorylated MtrB thus had a higher affinity for NarL and TcrX than for its cognate partner MtrA. For three other non-cognate RRs, namely KdpE, PhoP and TcrA, the affinities were 868±100 nM, 3580±121 nM, and 3071±74 nM, respectively, all lower than for MtrA. The order of affinities of phosphorylated MtrB for the different RRs tested was thus NarL>TcrX>MtrA>KdpE>TcrA>PhoP.

**Figure 1.**
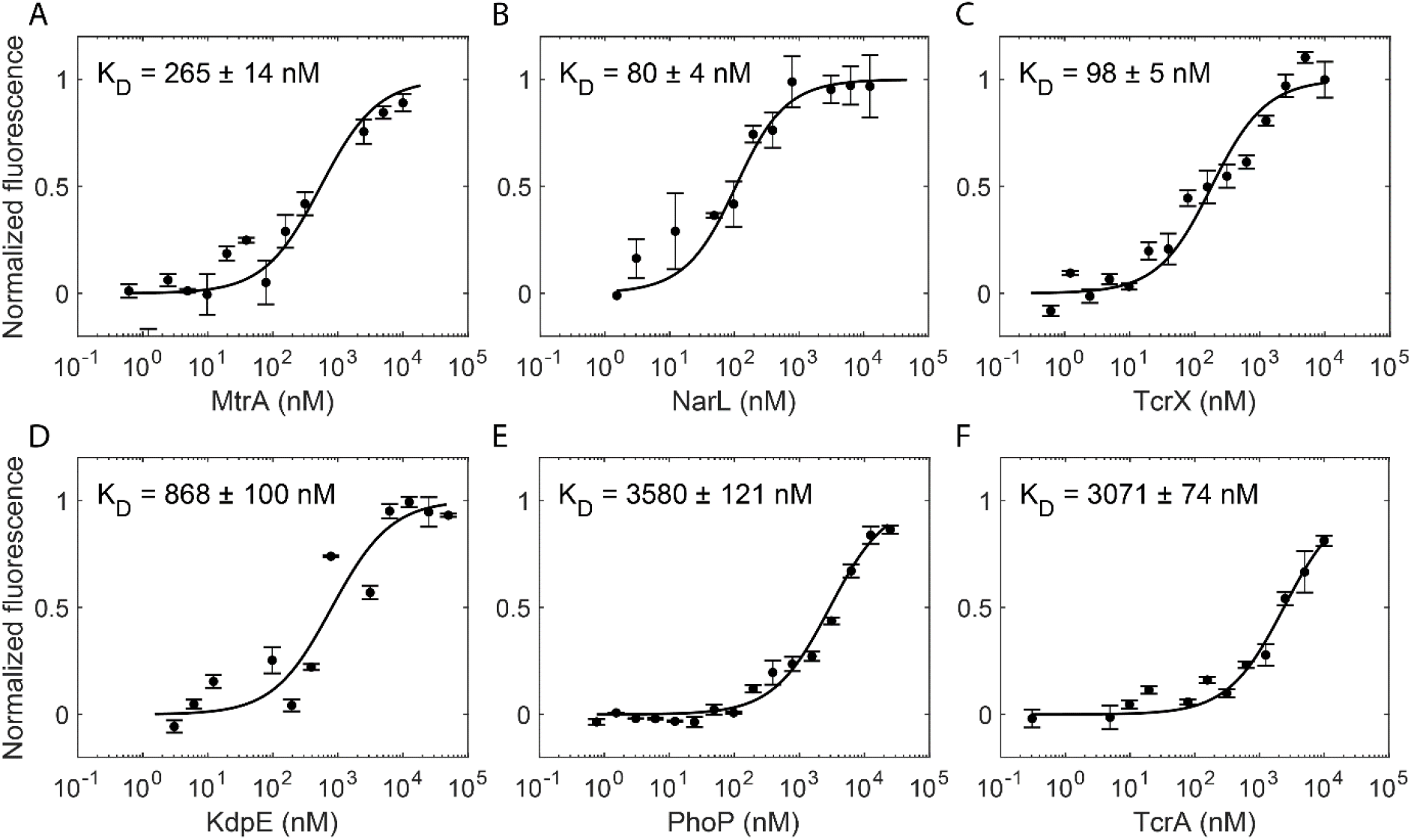
Binding affinities of phosphorylated MtrB for cognate and non-cognate RRs. Changes in the thermophoretic movement of 50 nM of fluorescently tagged MtrB post autophosphorylation, P∼MtrB-GFP, were measured as a function of titrant concentration for the titrant RRs (concentration range): **(A)** MtrA (0.61 nM to 10 μM), **(B)** NarL (1.5 nM to 12.5 μM), **(C)** TcrX (0.61 nM to 10 μM), **(D)** KdpE (3.1 nM to 50 μM), **(E)** PhoP (0.76 nM to 25 μM), and **(F)** TcrA (0.31 nM to 10 μM). K_D_ values were evaluated by plotting normalized fluorescence against the logarithmic concentrations of serially diluted ligand (RRs). Symbols are mean ± S.E.M. from more than 3 independent experiments and curves are best-fits.

In the unphosphorylated state, the binding affinities of MtrB were consistently lower (Figure S1). The *K*_*D*_ of GFP-tagged MtrB for MtrA was 2183±368 nM, indicating an over 8-fold weaker binding than its phosphorylated analog. Besides, the above rank ordering was not conserved in the unphosphorylated state. For the two non-cognate RRs we examined, namely, TcrX and PhoP, the affinities were 21715±5022 nM and 18087±2736 nM, respectively, so that the rank ordering was MtrA>PhoP>TcrX (Figure S1). While this is consistent with the expectation of specificity residues leading to better cognate HK-RR binding than non-cognate pairs^23^, phosphorylation of HK, in our studies, appeared to influence this rank ordering significantly. Signal transduction requires phosphorylated HKs, which we therefore focused on in our subsequent studies.

We first tested whether this preferential binding of phosphorylated HKs to non-cognate RRs was also seen with other TCSs of *M. tuberculosis*.

### Preferential binding of phosphorylated HKs to non-cognate RRs appears widely prevalent

We measured the binding affinities of four other phosphorylated, GFP-tagged (for MtrB and PhoR) and histidine tag labelled HKs of *M. tuberculosis* for their cognate as well as several non-cognate RRs, the latter implicated previously to be involved in crosstalk with the corresponding HKs^19^. Remarkably, we found that in every case there was at least one non-cognate RR for which the HK had a higher affinity than its cognate partner (Figure 2, Figure S2, Figure S3). The HK PhoR had a higher affinity for the non-cognate RRs TcrX and DevR (*K*_*D*_=484±82 nM and 154 ± 28 nM, respectively) than its cognate partner PhoP (*K*_*D*_=1485±203 nM) and another non-cognate RR TcrA (*K*_*D*_=1827±163 nM); the HK KdpD had a higher affinity (*K*_*D*_=355±44 nM) for NarL than its cognate partner KdpE (*K*_*D*_=4494±853 nM); the HK DevS had a higher affinity (*K*_*D*_=325±48 nM) for NarL than its cognate partner DevR (*K*_*D*_=13234±1663 nM); and the HK PrrB had a higher affinity (*K*_*D*_=1960±251 nM) for the RR MprA than its cognate partner PrrA (*K*_*D*_=4308±386 nM).

**Figure 2.**
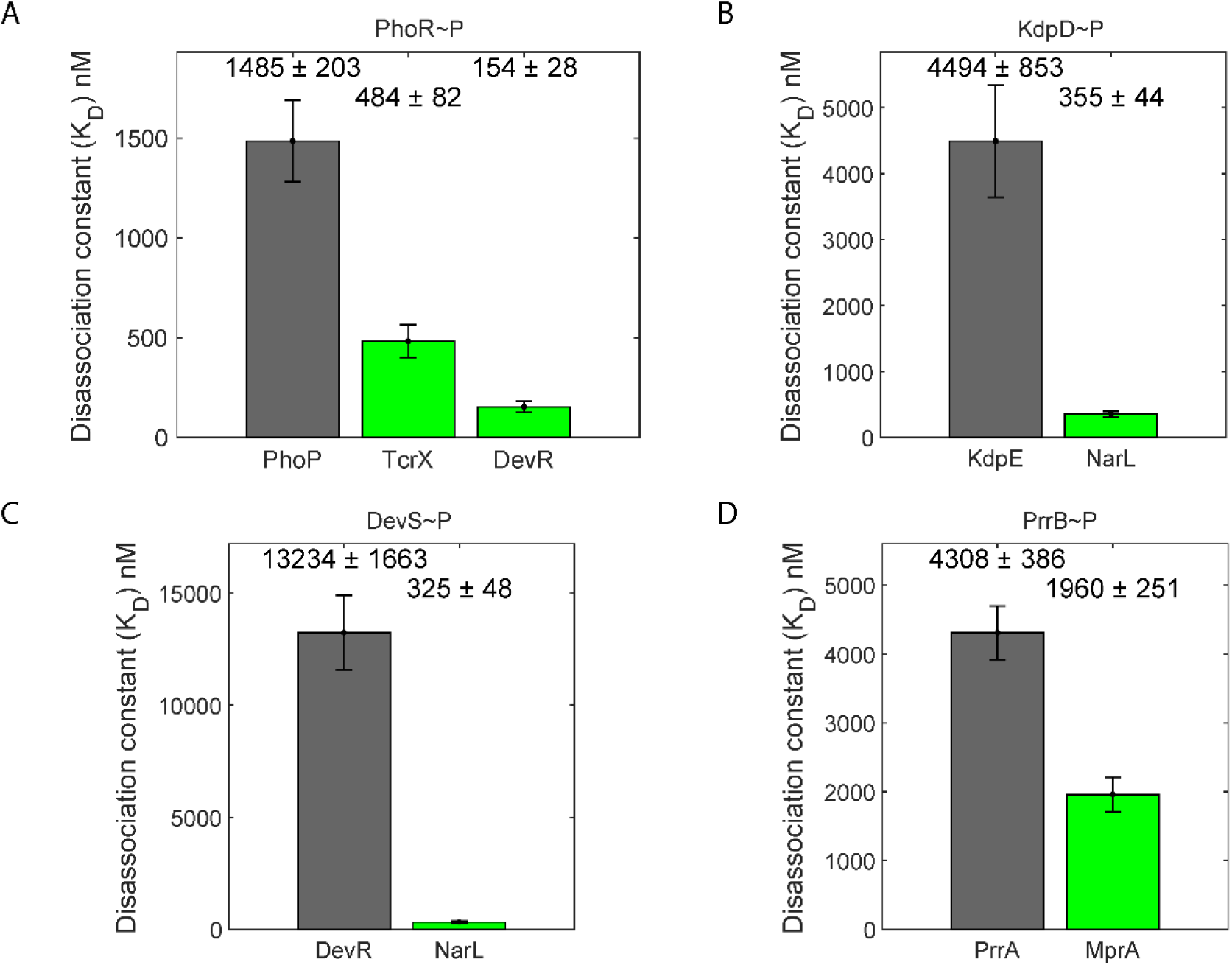
Binding affinities of several HKs for their cognate and non-cognate RRs. Affinities measured as in Figure 1 for phosphorylated **(A)** PhoR, **(B)** KdpD, **(C)** DevS, and **(D)** PrrB, for their respective cognate (grey) and some non-cognate (green) RRs implicated in crosstalk with the HKs. The affinities as mean ± S.E.M. from at least 3 repeats are indicated. Detailed measurements leading to the affinity estimates are in Figures S2 and S3, including for any non-cognate RRs with weaker affinities than the cognate ones.

We thus found that in every one of the five TCSs of *M. tuberculosis* we examined, including MtrAB, the phosphorylated HKs had a higher binding affinity for some non-cognate RR than their cognate partners. The affinities were at least 2-fold higher (for PrrAB) but could be over 40-fold higher (for DevRS) for the non-cognate RRs compared to the cognate partners. Phosphorylated MtrB, for instance, would thus bind preferentially to the non-cognate RRs NarL and TcrX over its cognate counterpart MtrA. Similarly, phosphorylated DevS would bind NarL in preference over DevR. Because phosphotransfer to non-cognate counterparts is typically inefficient^24,25^, this binding could amount to the sequestration of the HKs from their cognate RRs and the arrest of signal transduction through the cognate pathway. We examined next the impact of this preferential binding order on phosphotransfer and signal transduction. We focused on the MtrAB TCS.

### Sequestration of MtrB by the non-cognate RR NarL inhibits phosphotransfer to the cognate RR MtrA *in vitro*

We co-incubated the phosphorylated HK MtrB with its cognate RR, MtrA, in the presence or absence of the higher affinity non-cognate RR NarL tagged with mRuby, termed NarL-mRuby. We measured the levels of phosphorylated MtrB, MtrB∼P, and phosphorylated MtrA, MtrA∼P, as functions of time to assess the extent of phosphostransfer. The RRs were both in 2-fold excess (100 pmol each) of the HK (50 pmol), so that phosphotransfer was not limited by the availability of RRs. We found that as the MtrB∼P levels decreased, MtrA∼P levels rose (Figure 3A and B). In the presence of NarL, the latter rise was subdued. Whereas peak MtrA∼P levels reached 30% (a.u.) without NarL, they remained at 20% with NarL (Fig. 3D and E). In the absence of MtrA, the decline of MtrB∼P was far lower; whereas MtrB∼P levels declined to ∼20% of their initial value in 20 min in the presence of MtrA, they remained at ∼60% without MtrA (Figure 3C and F), ruling out significant phosphotransfer from MtrB∼P to NarL. In another control, we replaced NarL-mRuby with another RR PdtaR-RFP, which did not interact with MtrB, and found that peak MtrA∼P levels reached 30%, ruling out any non-specific effects of non-cognate RRs or their fluorescent tags on the phosphotransfer reaction (Figure S4). The reduced MtrA∼P levels with NarL were thus a consequence of the ‘sequestration’ of MtrB∼P by NarL.

**Figure 3.**
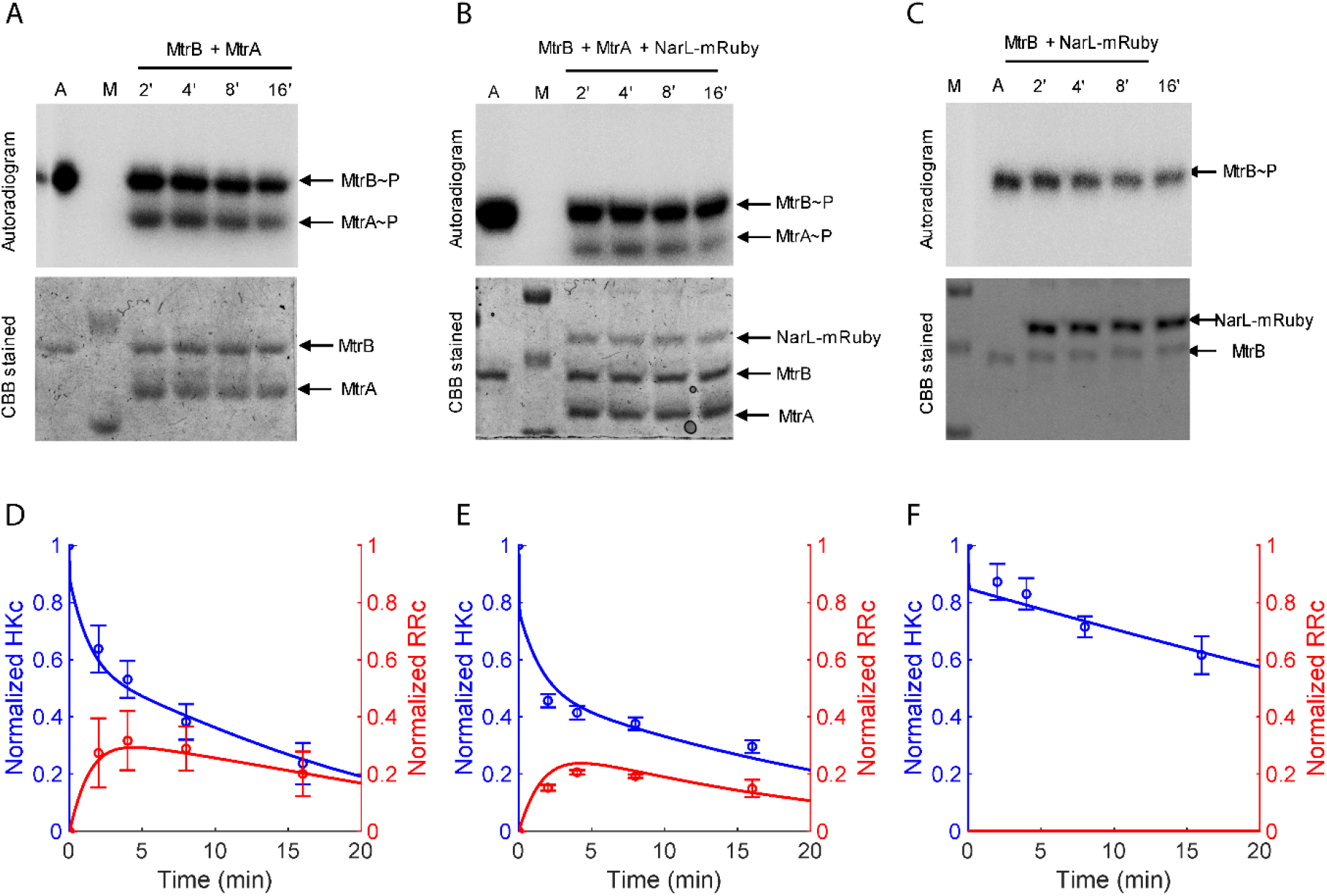
Phosphotransfer kinetics from MtrB to MtrA with and without sequestration. Time course assay of the phosphotransfer from the phosphorylated HK MtrB∼P to the cognate RR MtrA in the **(A)** absence or **(B)** presence of the non-cognate RR NarL-mRuby. **(C)** The same as (B) but in the absence of MtrA. M represents marker and A represents autophosphorylation control of MtrB∼P. Top panels in (A)-(C) are autoradiograms and bottom panels corresponding Coomassie Brilliant Blue (CBB) stained gels. **(D)-(F)** Densitometric analysis of the time course assays in (A)-(C), respectively. The autophosphorylation control was used to normalize the intensities of the individual bands. Blue symbols represent MtrB∼P and red symbols MtrA∼P. Lines represent best-fits of our model (Methods). The error bars represent mean ± S.E.M (n=4).

A consequence of the sequestration is the suppression of signaling through the cognate pathway. It would imply that higher levels of stimulation of the cognate pathway would be required to trigger positive autoregulation and mount a significant cognate response. To establish this concretely and to elucidate the design principle underlying the widely prevalent sequestration by non-cognate RRs we observed, we constructed a mathematical model.

### Mathematical model of TCS signaling in the presence of non-cognate RR

We developed a mathematical model of TCS signaling *in vivo* with positive autoregulation and the presence of a non-cognate RR (Figure 4; Methods).

**Figure 4.**
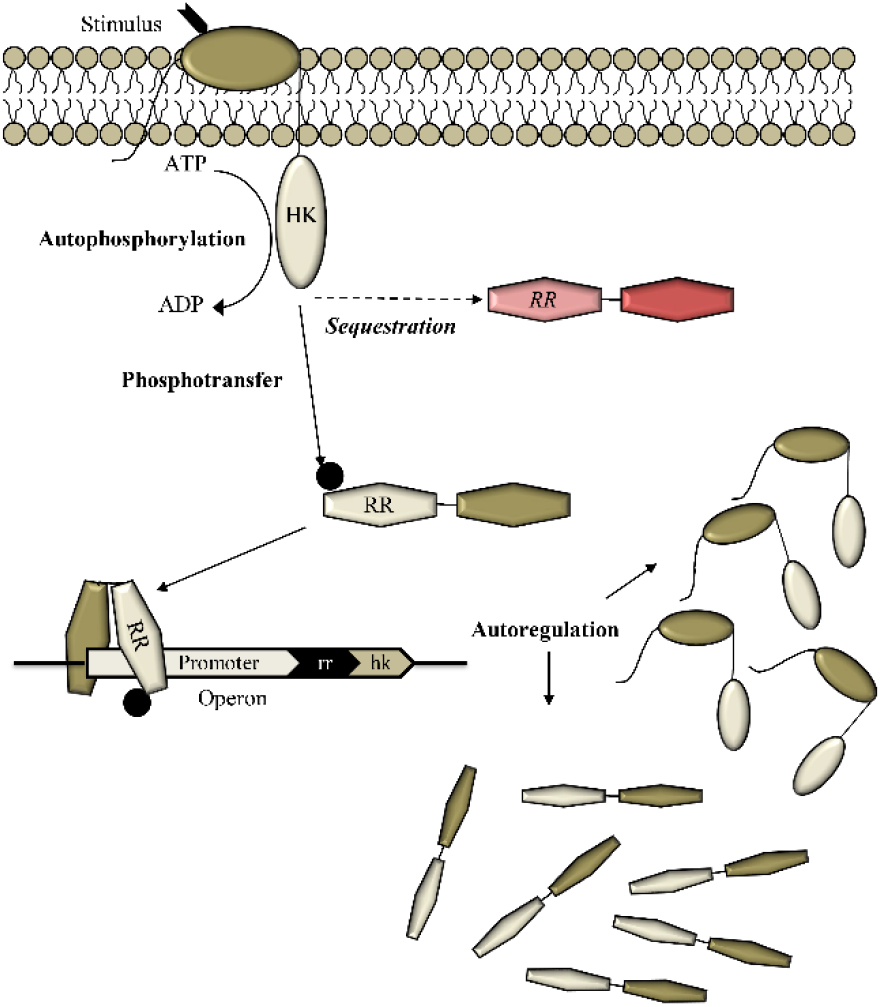
Schematic of the mathematical model. The model considers an extracellular stimulus triggering the autophosphorylation of HK and activating a TCS pathway. The phosphorylated HK could transfer phosphoryl group to its cognate RR, which could bind DNA and trigger the synthesis of the HK and RR proteins, marking positive autoregulation of the TCS. The phosphorylated HK could bind non-cognate RRs (red) preferentially, when the latter have higher affinity for the HK than its cognate counterpart, resulting in HK sequestration and the suppression of cognate signaling. Only with a sufficiently strong stimulus does sufficient HK autophosphorylation result leading to cognate RR binding despite sequestration and the mounting of a response through the cognate pathway. Equations (1)-(15) list the reaction events in this model. The rate equations are in Methods.

The following events comprise the model.

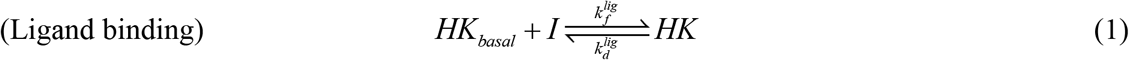

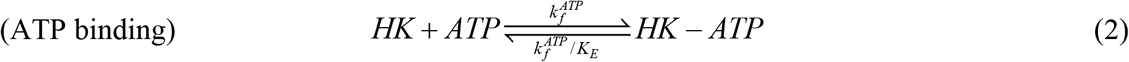

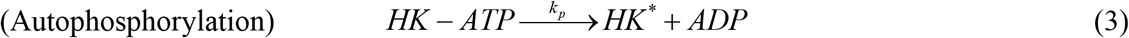

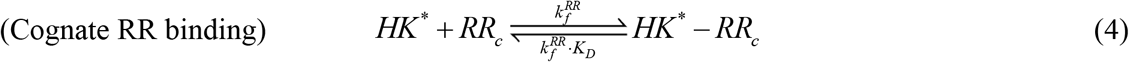

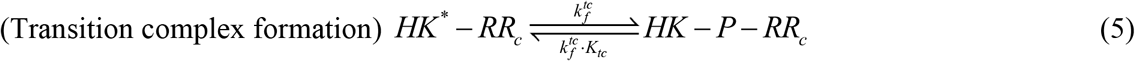

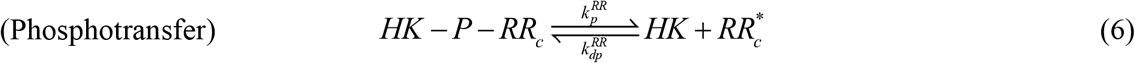

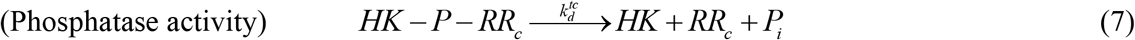

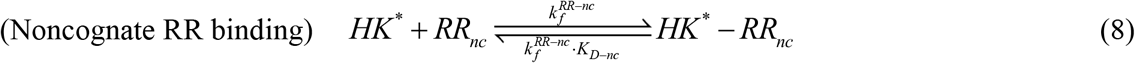

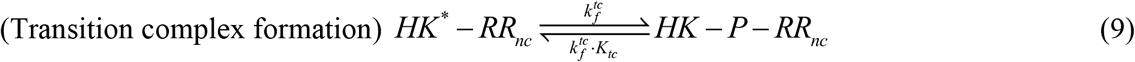

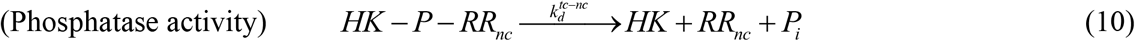

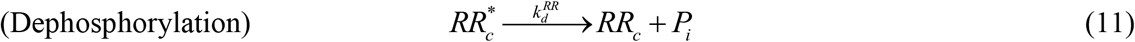

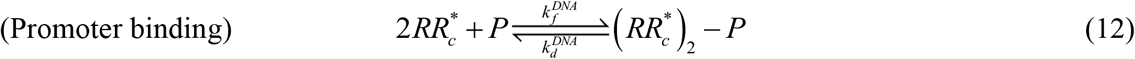

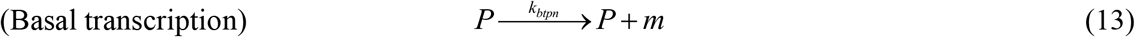

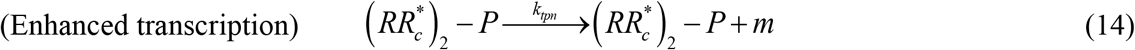

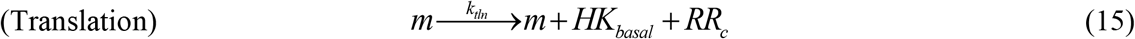

Briefly, the model considers HK in a basal state, which is activated by an input or stimulus, *I*, (Eq. 1), following which the HK binds ATP (Eq. 2) and gets autophosphorylated (Eq. 3). The phosphorylated HK, denoted HK^*^, binds the cognate RR, denoted RR_c_ (Eq. 4), and forms a transition complex, HK-P-RR_c_, poised for phosphotransfer (Eq. 5). The complex either effects phosphotransfer (Eq. 6) or dissociates with the loss of the phosphoryl group into inorganic phosphate, *P*_*i*_, the latter consistent with the phosphatase activity of HK (Eq. 7). In the presence of the non-cognate RR, denoted RR_nc_, the phosphorylated HK can bind RR_nc_ (Eq. 8), form a transition complex (Eq. 9), and get dephosphorylated (Eq. 10). The transition complex here is assumed not to be able to effect phosphotransfer to RR_nc_. The phosphorylated RR_c_, denoted RR_c*_, can get dephosphorylated spontaneously (Eq. 11), or dimerize and bind the promoter region P (Eq. 12). The basal transcription of downstream genes (Eq. 13) yielding mRNA, *m*, now happens at an enhanced rate (Eq. 14). Translation of the mRNA results in the production of the HK and RR_c_ (Eq. 15), closing the positive autoregulation loop, and other response proteins.

We constructed rate expressions for the above events, which resulted in a set of coupled ordinary differential equations (Methods). The parameters (rate constants) and their estimated values are listed in Table S1. Most parameters were set to values known from previous studies. We estimated the remaining parameters by fitting model predictions excluding the autoregulatory loop to the *in vitro* data above.

### Mathematical model fits *in vitro* data

We applied our reduced model (Eqs. (2)-(11)) using the corresponding rate equations (Methods) to fit time-courses of MtrB∼P and MtrA∼P with and without NarL (Figure 3). We also simultaneously fit data of MtrB∼P levels in the control experiment without MtrA. The model provided good fits to the data (Figure 3), reiterating the observation that sequestration by non-cognate RRs can suppress the phosphorylation of the cognate RR. The best-fit parameters estimated are in Table S1. With all the parameters thus identified, we applied the full model to examine how the sequestration would influence signal transduction via the cognate TCS pathway *in vivo*.

### Sequestration by non-cognate RRs establishes a threshold for cognate TCS signal transduction

We considered the input, *I*, to represent an environmental cue that rises sharply and then declines exponentially: *I*=*I*_0_exp(-*t/τ*). Here, *I*_0_ represents the strength of the signal and *τ* its duration. A large *I*_0_ and a small *τ* would represent a strong but fleeting stimulus, whereas a small *I*_0_ and a large *τ* would be a weak but lasting stimulus (Figure 5A). We solved our model for a wide range of values of *I*_0_ and *τ* and examined the response. We quantified the response, or output, as the fractional occupation of the promoter region by the cognate RR (Methods).

**Figure 5.**
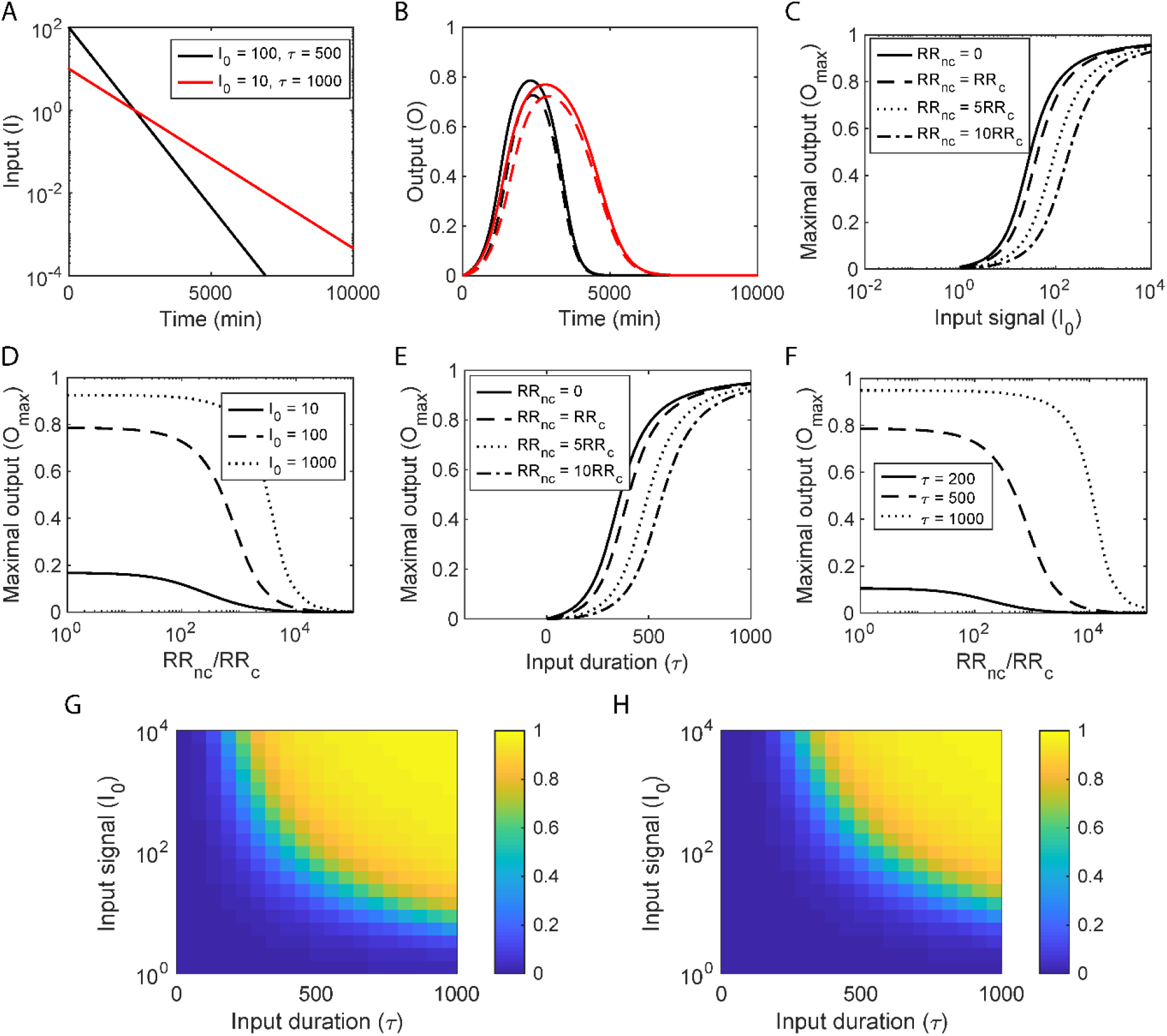
Model predictions of TCS signal transduction and the impact of sequestration. **(A)** Representative inputs, *I*, indicating strong but short-lived (black) and weak but extended (red) stimuli. **(B)** The corresponding outputs without (solid lines) and with (dashed lines) sequestration by a non-cognate RR. **(C)** The peak of the response (*O*_*max*_) as a function of the maximum input, *I*_0_, for different extents of sequestration, determined by the ratios of the non-cognate RRs, RR_nc_ to the cognate RR, RR_c_, indicated. **(D)** *O*_*max*_ as a function of the ratio RR_nc_/RR_c_ for different *I*_0_. **(E)** *O*_*max*_ as a function of the signal half-life, *τ*, for different values of RR_nc_/RR_c_. **(F)** *O*_*max*_ as a function of RR_nc_/RR_c_ for different values of *τ*. (*τ* is in minutes throughout.) Heatmaps showing *O*_*max*_ as functions of *I*_0_ and *τ* in the **(G)** absence or **(H)** presence of non-cognate RRs, indicating the threshold stimulation for response shifting to higher *I*_0_ and *τ* with sequestration. Corresponding calculations for the total response, *O*_*total*_, are in Figure S5. Model predictions were obtained by solving Eqs. (1)-(33) (Methods) using parameter values listed in Table S1.

For a given signal, the output, *O*, in the absence of sequestration ([RR_nc_]=0) rose, attained a maximum, *O*_max_, and then declined to zero (Figure 5B). (The output would stay elevated in response to a persistent signal (*τ*→∞); our focus here was on temporary and weak signals.) Increasing *I*_0_ or *τ* increased the maximum response, indicating that stronger or more sustained stimuli led to stronger responses. In the presence of sequestration ([RR_nc_]>0), however, the rise in the output was delayed, the maximum output was suppressed, and the output vanished sooner (Figure 5B). In effect, sequestration by non-cognate RRs suppressed signal transduction through the cognate pathway.

We examined next how the maximum output, *O*_*max*_, and the cumulative output, *O*_*total*_, (area under the output-time curve) varied with *I*_0_ and *τ* in the absence and presence of sequestration. We found that *O*_*max*_ exhibited a sigmoidal dependence on *I*_0_, remaining low until a threshold level, *I*_*threshold*_, was crossed, rising sharply thereafter, and then reaching a saturation level, *O*_*sat*_, where further increases in *I*_0_ triggered marginal increases in *O*_*max*_ (Figure 5C). Such nonlinear, sigmoidal responses are attributed to positive autoregulation.^5^ With sequestration, we found that the sigmoid shifted to the right, i.e., to higher values of *I*_0_. Thus, in particular, *I*_*threshold*_ increased, indicating that a much larger level of stimulation was necessary for a significant response to be mounted. The shift increased with the level of RR_nc_, amounting to greater sequestration.

Conversely, for a given level of stimulation, *I*_0_, the maximum response, *O*_*max*_, exhibited an inverse sigmoidal dependence on RR_nc_ (Figure 5D). As RR_nc_ increased from zero, *O*_*max*_ decreased gently until a critical RR_nc_ was crossed. At this point, *O*_*max*_ decreased sharply and reached minimal levels. The critical RR_nc_ was the value for which *I*_0_ became comparable to the threshold stimulation level, *I*_*threshold*_. Any RR_nc_ above this would amount to sequestration being strong enough that *I*_0_ would remain below *I*_*threshold*_, yielding a weak response. Thus, sequestration prevents the mounting of a strong response to short-lived stimuli.

Similar behavior was observed when *I*_0_ was kept constant and *τ* was varied. As *τ* increased, *O*_*max*_ rose in a sigmoidal manner, indicating the existence of a threshold duration, *τ*_*threshold*_, below which the output was weak and above which the output rose sharply to saturation, *O*_*sat*_ (Figure 5E). As the level of sequestration increased, i.e., as RR_nc_ rose, *τ*_*threshold*_ increased, indicating that the stimulus had to last longer for a significant response to be mounted (Figure 5D). Again, for a given *τ, O*_*max*_ exhibited an inverse sigmoidal dependence on RR_nc_, indicating the existence of a critical RR_nc_ at which *τ* became comparable to *τ*_*threshold*_ and above which *τ* was unable to elicit a significant response (Figure 5F).

Heat maps comparing *O*_*max*_ as a function of *I*_0_ and *τ* with (RR_nc_>0) and without (RR_nc_=0) sequestration show that *O*_*max*_ was suppressed by sequestration when *I*_0_, *τ* or both were small (Figure 5G and H). The same trend applied to *O*_*total*_ (Figure S5). Sequestration thus appeared as a design to prevent the mounting of a strong response despite positive autoregulation when the stimulation was weak or fleeting.

With this insight from mathematical modeling, together with evidence from our *in vitro* experiments, we examined whether this design was evident *in vivo* using our experimental models.

### Non-cognate RR suppresses TCS signal transduction *in vivo*

Our *in vitro* experiments above demonstrated the sequestration of MtrB∼P by NarL. We examined the effect of this sequestration *in vivo* using *Mycobacterium bovis* BCG, a non-pathogenic surrogate for *M. tuberculosis*. The stimulus for the MtrAB system is unknown.^26^ The system is known to be active during cell proliferation and regulates the expression of the downstream gene *dnaA*.^27^ Because the stimulus is unknown, tuning it to define threshold responses was not possible. However, the expression level of NarL, a higher affinity non-cognate RR for MtrB, could be tuned, enabling control of the level of sequestration. We increased the level of NarL using the anhydrotetracycline (aTC) tunable expression system (pTIC6)^28^ and measured the changes in the transcript level of the gene *dnaA* as a function of NarL expression. Supporting our *in vitro* data and the expectation from our mathematical model, about a 5-fold increase in NarL expression (increasing RR_nc_) caused a nearly 35% reduction in the *dnaA* transcript level, or output from the cognate pathway (Figure 6). This marked reduction in the level of *dnaA* due to the upregulation of the non-cognate RR, NarL, indicates the presence of the sequestration motif *in vivo*. Sequestration of phosphorylated HK by non-cognate RRs thus appears to be a design to prevent disproportionately amplified responses to weak or short-lived stimuli.

**Figure 6.**
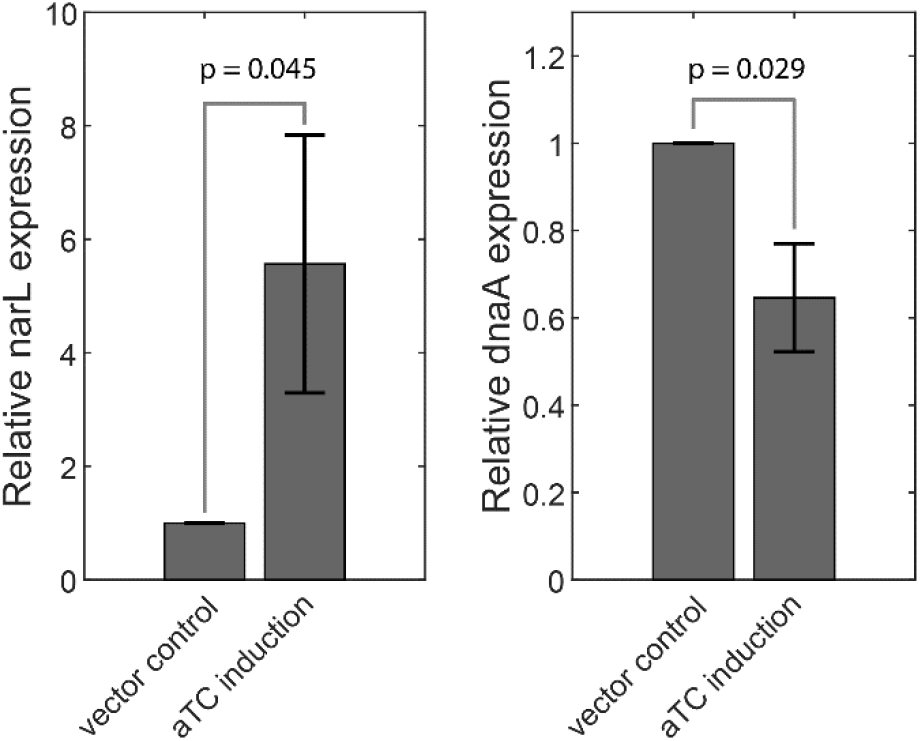
Effect of HK sequestration on the cognate TCS response *in vivo*. **(Left)** NarL expression in *M. bovis* BCG strains showing control levels and overexpressed levels achieved through a tetracycline inducible promoter. **(Right)** Corresponding expression levels of the gene *dnaA*, a downstream target of the MtrAB TCS. The gene expression was normalized to the expression levels of 16s rRNA, followed by the expression levels in the strain carrying only the vector (vector control). Data represent n=3 biologically independent experiments. Effect of HK sequestration is quantitatively observed as about 40% reduction in the output of the cognate TCS pathway due to the approximately five-fold increase in the non-cognate RR NarL expression.

## DISCUSSION

Our study reveals a new design principle underlying the regulation of bacterial two-component signaling (TCS) systems. Positive autoregulation of TCSs, where a TCS upregulates its own proteins in response to stimulation, is a widely prevalent feature that aids signal amplification and the mounting of a strong and lasting response to stimuli.^8^ A downside of positive autoregulation, however, is that a disproportionate response may be triggered to weak or fleeting stimuli, which may cost resources and hence reduce bacterial fitness. How bacteria overcome this limitation has been puzzling. Here, we unraveled a new mechanism, sequestration of HKs by non-cognate RRs, that provides an answer. We found with every one of the five mycobacterial TCSs we studied that HKs bind to at least one non-cognate RR more tightly than their cognate counterparts. The tighter-binding non-cognate RR thus sequesters the HK until a strong enough stimulus results in sufficient HK autophosphorylation that a ‘leak’ to the cognate RR ensues. Positive autoregulation then amplifies the HK and cognate RR protein levels, allowing significant signaling through the cognate pathway and the mounting of a proportionate response. We demonstrated this design principle using mathematical modeling as well as *in vitro* and *in vivo* experiments on mycobacterial TCSs.

The costs and benefits of positive autoregulation have been investigated extensively because of their importance to the regulation and control of biological processes in diverse settings.^5,7,29-31^ Among the costs identified has been the delay in the response to a stimulus arising from the need to build up a sufficient amount of the entities involved before positive autoregulation can begin to amplify them.^29-31^ When the positive autoregulatory network is made more sensitive, it allows a gain in speed, but could lead to excessive amplification of the long-term response, resulting in a substantial fitness cost. A coupled negative feedback loop has been proposed recently as a way to ensure speed in the response without the excessive long-term protein production.^30^ Here, we recognize yet another limitation of positive autoregulation. The positive autoregulation with a coupled negative feedback describes how a fast and controlled response, which is appropriate, can be mounted to a stimulus that is substantial and lasting. The sensitive positive autoregulation ensures speed, whereas the coupled negative feedback prevents uncontrolled amplification. When the stimulus is weak or fleeting, however, for which a response may not be warranted, let alone a speedy one, the sensitive positive autoregulation may still initiate a response and amplify it, introducing a fitness cost. In the context of bacterial TCSs, we showed here that sequestration of HKs by non-cognate RRs introduces a threshold level of stimulation for a response to be mounted and prevents such unnecessarily speedy and amplified responses. Thus, while the negative regulation controls the response in the late stages, the sequestration we identified controls it in the early stages of stimulation. Together, the two present a more complete regulatory design of bacterial TCSs under positive autoregulation.

Mycobacterial TCSs have been found to engage in extensive cross-talk, where HKs transfer phosphoryl groups to non-cognate RRs.^19^ A pre-requisite for this cross-talk is the binding of phosphorylated HKs with non-cognate RRs. The binding affinities of these interactions, however, had not been measured thus far. Here, we employed the novel, facile, solution-based technique, microscale thermophoresis^32,33^, to estimate the binding affinities of several cognate and non-cognate HK-RR pairs from within the sets that were shown to cross-talk *in vitro*. Remarkably, we found for every HK the presence of at least one, but often multiple, non-cognate RRs that had higher affinity than the cognate counterpart. This finding, together with the recognition that phosphotransfer to non-cognate RRs is slow and inefficient compared to cognate RRs^25^, provided the basis for sequestration as a regulatory design principle that we unraveled.

The extent of regulation by this process would depend on the relative binding affinities as well as the expression levels of the cognate and non-cognate RRs. In our *in vivo* studies, we demonstrated that increasing the levels of the non-cognate RR NarL reduced the output of the cognate MtrAB TCS. This suggests that the sequestration motif may be prevalent even with non-cognate RRs that have lower binding affinities than the cognate ones provided their expression levels are sufficiently high. In effect, one may view the collection of all non-cognate RRs as the source of sequestration. The relative abundance of RRs is greater than their cognate HKs.^34-37^ A typical bacterial system may contain many tens to hundreds of distinct TCSs, thus providing a large pool of non-cognate RRs for any HK.^38^ Even if no single non-cognate RR were to have a higher binding affinity for an HK than its cognate counterpart, the total concentration of the non-cognate RRs may be sufficiently large that sequestration would result. Also, “orphan” RRs, for which cognate HKs are unidentified^26^, appear compelling candidates for functioning as sequesters. It is therefore, together with our observation of higher affinity non-cognate RRs for every HK we studied, that we speculate that sequestration is likely to be a widely prevalent design for ensuring that responses are mounted only after a threshold level of stimulation is realized.

We distinguish sequestration from sinks, the latter also argued to give rise to thresholds^18,22^. If non-cognate binding were to strip the HK of its phosphoryl group, then the non-cognate RR would act as a sink. It would become available for repeating the act with the next phosphorylated HK, creating a perpetual drain, or a sink, for phosphoryl groups. It is apparent that a sink would place a much harsher demand on the signal for eliciting a response because it would eliminate the possibility of signaling from any HK that it would bind. Sequestration, on the other hand, is likely to exert gentler control. It suppresses the signal temporarily and allows the HK to dissociate from the non-cognate RR with its phosphoryl group and retain its ability to trigger signaling through the cognate pathway. The circumstances that confer an evolutionary advantage upon sequestration over sinks remain to be elucidated.

In settings where sequestration is due to non-cognate RRs that bind more strongly to an HK than its cognate counterpart, targeting the non-cognate RR-HK binding may be a novel intervention strategy. It may compromise the ability of the bacterium to withhold responses to frivolous stimuli, draining its resources and reducing its viability. Further, the high affinity binding between the HK and non-cognate RR suggests the existence of specific binding regions, which could be targeted by drugs or vaccines.

In summary, our study presents evidence of a new paradigm of TCS signal regulation, where sequestration by non-cognate RRs ensures that responses are mounted only after a threshold level of stimulation is realized.

## METHODS

### Mathematical model of TCS signalling with sequestration of HK by non-cognate RR

Our model describes the signal transduction via TCS in the presence of non-cognate RR (Equations 1-15). We wrote the following rate equations to describe the dynamics of signal transduction following stimulation by ligand, *I*. The equations build on earlier models of TCS signaling^19,39^,40 and advance them by incorporating the role of non-cognate RRs.

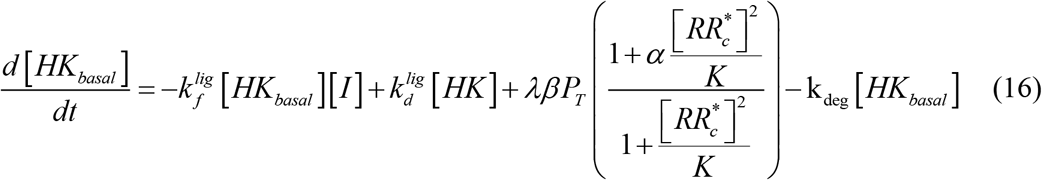

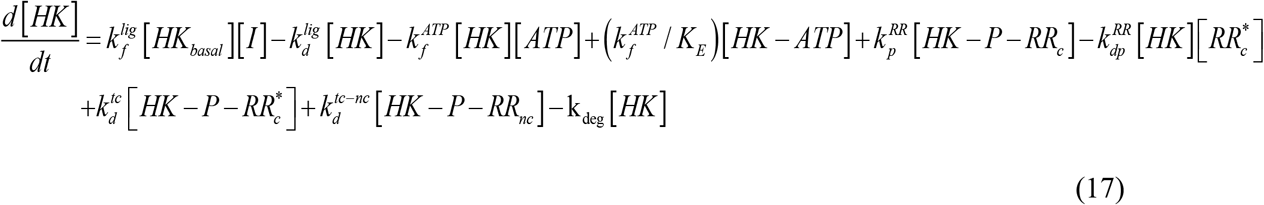

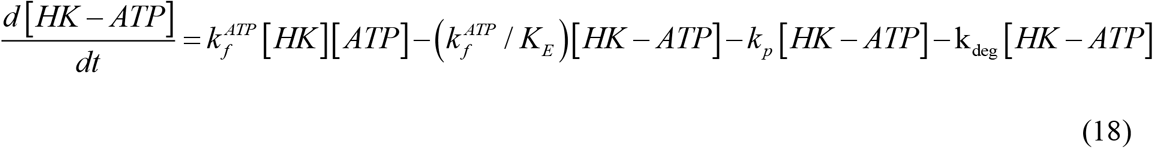

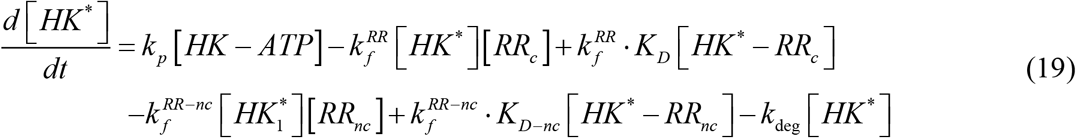

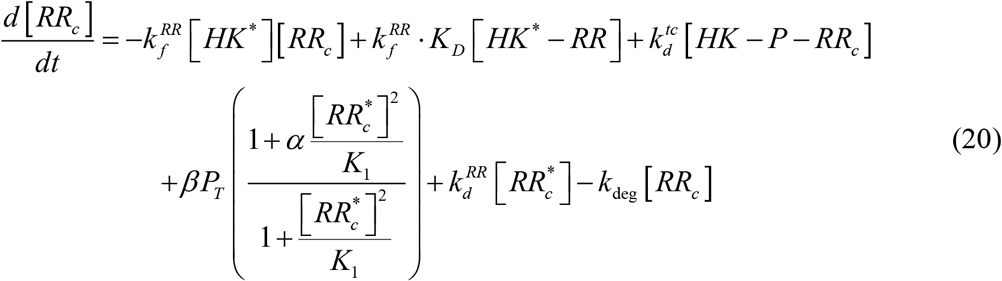

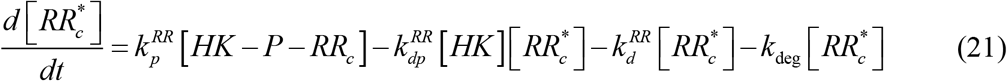

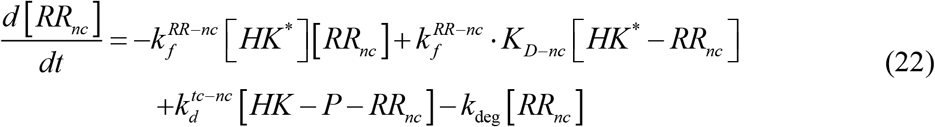

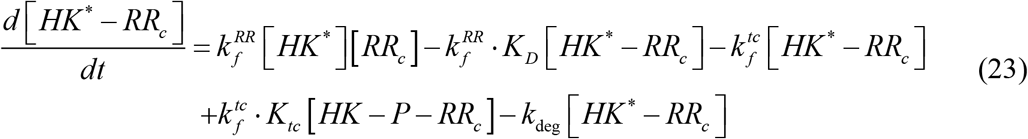

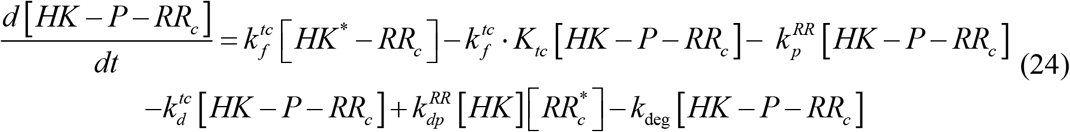

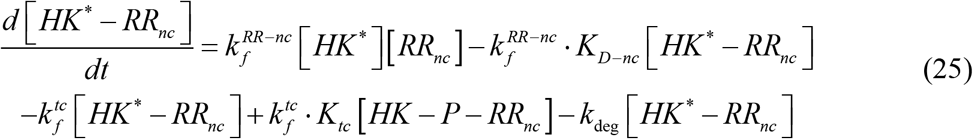

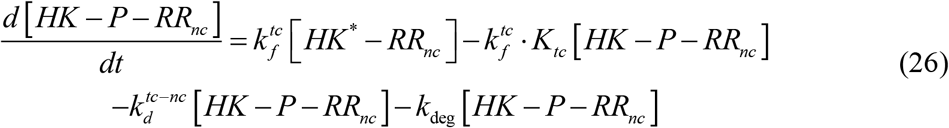

The equations were constructed by writing standard rate expressions for the events in Eqs. (1)-(15). Thus, the concentration of basal HK can change due to the following events: ligand binding and unbinding, autoregulation, and degradation. These form the four terms on the right-hand side of Eq. (16) above. Similar expressions were formulated for the other species (Eqs. 17-26). The meanings of the rate constants are in Table S1. We explain the non-obvious terms here. Following *HK*^*^ binding to *RR*_*c*_, the resulting complex,

*HK*^*^ − *RR*_*c*_, is assumed to form the transition complex *HK* − *P* − *RR*_*c*_, poised for phosphotransfer to *RR*_*c*_. A similar model, where the phosphoryl group is transferred within the complex, has been used earlier.^4^ The transition complex, *HK* − *P* − *RR*_*c*_, can either effect phosphotransfer yielding *HK* and 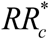 or dissociate, releasing inorganic phosphate (*P*_*i*_) and leaving behind unphosphorylated *HK* and *RR*_*c*_. This formalism is different from the classical phosphatase activity found in other models^19,39^, which ignore the transition complex. We considered the complex because of its importance to non-cognate RR binding. The non-cognate RR, RR_nc_, could form the analogous transition complex, *HK* − *P* − *RR*_*nc*_, but the latter complex was assumed not to be able to effect phosphotransfer. Because phosphatase activity is generally slower than phosphotransfer, the latter complex becomes relatively long-lived and acts as a sequester.

The autoregulation leading to the expression of HK and RR proteins was modeled using the pseudo-equilibrium approximation following earlier studies^19,40^. We let *P*_*T*_ be the total concentration of promoter binding sites present on the bacterial genome. If *f*_*b*_ and *f* _*f*_ were the fractions of promoter sites in the bound and free states respectively, then the equilibrium of the events in Eq. 12 would yield

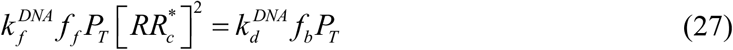

If we let 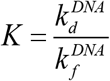 be the corresponding equilibrium dissociation coefficient, and since *f* _*f*_ + *f*_*b*_ =1, it followed that

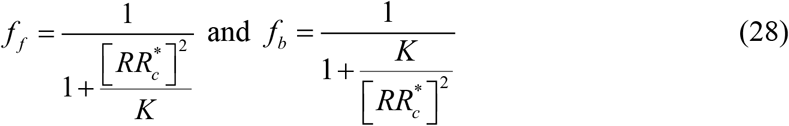

The change of mRNA concentration can be written from Eqns. 14 and 15 as

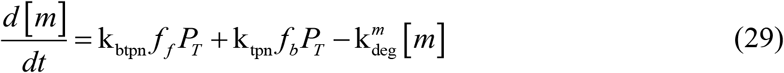

Applying the pseudo-equilibrium approximation^19,40^ for mRNA dynamics, i.e., 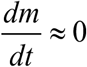, and using Eq. (28), we obtained

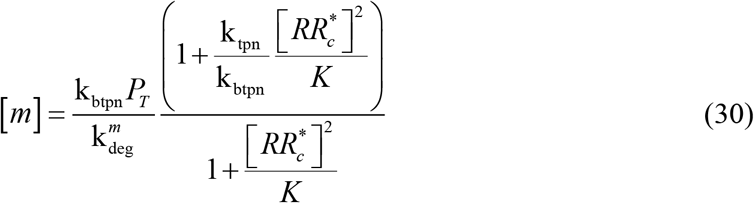

Translation of the mRNA molecules results in the production of the two TCS proteins *HK*_*basal*_ and *RR* molecules, with λ the ratio of the two production rates^19,40^.

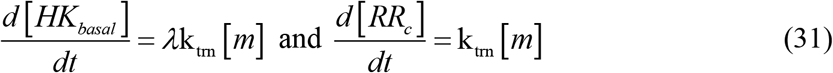

Substituting 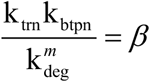 and 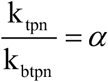, we obtained the synthesis rates of *HK*_*basal*_ and *RR*_*c*_ by translation as

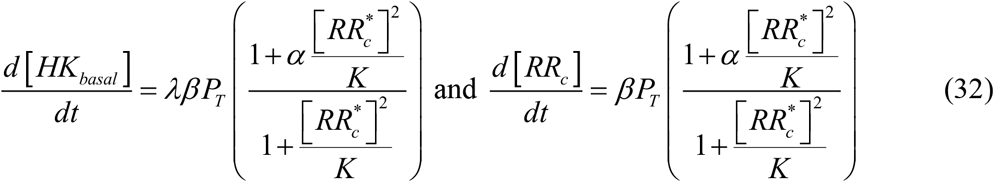

which we used in Eqs. (16) and (20) above to describe autoregulation. We defined the output, *O*, of the signal transduction events as the fraction of bound promoter regions, *f*_*b*_, so that:

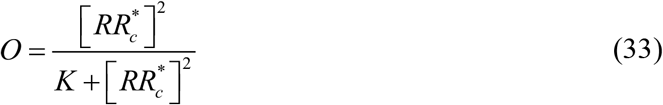

### Data fitting and parameter estimation

The parameter values employed are listed in Table S1 along with their sources. The dissociation constants between the cognate pairs MtrB and MtrA, *K*_*D*_, and non-cognate pairs MtrB and NarL, *K*_*D*−*nc*_, were estimated using microscale thermophoresis (Figure 1). The autophosphorylation of MtrB has been studied earlier and the equilibrium constant, *K*_*E*_, and the phosphorylation rate, *k*_*p*_, estimated.^41^ *ϕ*_*HK*_, the fraction of MtrB active was also estimated.^41^ The natural dephosphorylation rate of 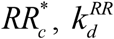 has been estimated for PhoB.^42^ We assumed the same natural dephosphorylation rate for the other RRs. The activation and deactivation rates for HKs, 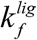 and 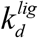, the forward rates for *HK* binding to 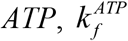, and of *HK*^*\**^ binding to *RR*_c_ and 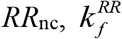 and 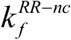, respectively, as well as the total promoter concentration, *P*_*T*_, were chosen from an earlier study.^39^ The equilibrium dissociation constant for DNA binding, *K*, the transcription rate, *α*, the translation rate, *β*, and the ratio of HK and RR produced by translation, λ, were from another previous study.^40^

The unknown parameters were the percentage activity of RR_c_, *ϕ*_*RR*_, the equilibrium dissociation constant of the transition complex, *K*_*tc*_, the phospho-transfer rate constant between MtrB and 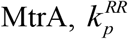, the binding rate constant of *HK* and 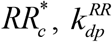, the phosphatase activity rate constant between MtrB and MtrA, 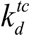, and the phosphatase activity rate constant between MtrB and NarL, 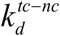. We assumed that the phosphotransfer rate from MtrB to NarL was negligible and set it to zero. The forward rate of transition complex formation, 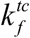, was assumed to be 100 min^-1^, to represent fast dynamics typically associated with transition complexes compared to other processes.

To estimate the unknown parameters, we simultaneously fit our model to data from the three time course assays in Figure 3. The termination of reactions resulted in denaturing of protein complexes. Therefore, while measuring *HK* ^*^ after reaction termination, we assumed that the complexes *HK*^*^ − *RR*_*c*_ and *HK*^*^ − *RR*_*c*_ could denature to *HK* ^*^ so that the concentration which we measured at each time point was the sum of *HK* ^*^, *HK*^*^ − *RR*_*c*_ and *HK*^*^ − *RR*_*nc*_ species. All the concentrations were normalized to the initial total active HK, *ϕ*_*HK*_. We solved the model using the MULTISTART function in MATLAB to do a global parameter search and used the LSQCURVEFIT function in MATLAB for optimization. The model equations were solved using ODE23 in MATLAB.

### Experimental Methods

#### Materials

All the media chemicals, biochemicals, and protein reagents were purchased from Sigma Merck (USA); antibiotics and DTT (Dithiothreitol) from Goldbio (USA); protein markers were from Thermo Fisher (USA), agarose-GSH resin and Ni^+2^-NTA resin were from GE Healthcare (USA); and restriction enzymes from Thermo Fisher (USA). Primers were synthesized from Bioserve (India) and *γ*^32^P ATP (>3500 Ci/mmol) was purchased from BRIT-Jonaki (India).

#### Bacterial strains and plasmids

Protein overexpression was carried out in *E. coli* Origami and Origami B (Novagen Inc., USA). These strains carrying the recombinant plasmids were propagated in LB containing ampicillin (100 μg/ml). The recombinant plasmids used for protein overexpression have been reported previously^19^. In brief, for the HKs, the plasmids containing only the cytosolic catalytic domains were used and for RRs the entire coding gene was used. For all GFP (Green Fluorescent Protein) tagged proteins, the GFP gene was cloned downstream of cytosolic catalytic domain of the respective HK with a linker region which encoded for the GSGGG spacer peptide, which facilitated functional separation of the two proteins as reported previously^22^.

#### Recombinant protein purification

For recombinant protein overexpression and purification, *E. coli* strains carrying the recombinant plasmid were grown at 37°C, in 200 ml of 2x YT broth until OD600 of 0.4 - 0.6, then IPTG (0.5 mM) was added to the culture and incubated further for 15-20 h at 12-15°C for protein overexpression. Cells were harvested by centrifugation and stored until use at -80°C. For protein purification in soluble conditions, the protocol described previously was followed^22^.

#### Autophosphorylation and phosphotransfer activity of GFP-tagged HKs using PAGE/autoradiography

The functional validation of the activity of GFP tagged HKs was conducted as previously reported for MtrB-GFP^27^. In brief, 5 μM of the purified GFP tagged HK, i.e., PhoR-GFP was incubated in the kinase buffer (50 mM Tris-HCl, pH 8.0, 50 mM KCl, 10 mM MgCl_2_) containing 50 μM ATP and 1μCi of γ^32^P-labelled ATP at 30°C for 2 hours (Figure S2A). Equimolar amount of the recombinant cognate RR PhoP diluted in kinase buffer was added to allow phosphotransfer for 5 minutes. The reaction was terminated by adding 1× SDS-PAGE sample buffer. The samples were resolved on 12% v/v SDS-PAGE. After electrophoresis, the gel was washed and exposed to phosphor screen (Fujifilm Bas cassette2, Japan) for 4 hours followed by imaging with Typhoon 9210 phosphorimager (GE Healthcare, USA).

#### Determination of affinities of HK∼P for RR using microscale thermophoresis (MST)

5 μM of the purified GFP-tagged HK was autophosphorylated as mentioned above using 50 μM ATP at 30°C for 90 min. The autophosphorylated HK∼P (50 nM) was mixed with increasing concentrations of titrant RRs (concentration range mentioned in figure legends), in the autophosphorylation buffer and kept at 30°C for 5 min. The sample was then loaded into standard treated capillaries and analyzed using a Monolith NT.115 (NanoTemper Technologies Germany). The blue laser was used for a duration of 35 s for excitation (MST power = 60%, LED power 40%). For the interactions involving fluorescently labelled recombinant proteins, the HKs were autophosphorylated and then mixed with 50 nM of Monolith NT His-tag labelling Kit Red Tris-NTA (L008; nanotemper Technologies Germany) incubated at 30°C for 30 minutes to allow labelling of the His-tag with the red fluorescent dye NT-647 and the red laser was used for a duration of 35 s for excitation (MST power = 60%, LED power 90%). The data were analyzed using MO Control software (NanoTemper Technologies Germany) to determine the K_D_ for the interacting proteins.

#### Time course analysis of phosphotransfer assays

In the phosphotransfer assays, 100 pmoles of the RRs diluted in kinase buffer were added to the reaction containing 50 pmoles of autophosphorylated HK, followed by incubation at 30°C for indicated time points. The reaction was terminated after the incubation by adding 1x SDS-PAGE gel loading buffer and resolved on a 15% SDS-PAGE gel. After electrophoresis, the gels are washed with deionized water and exposed on a phosphorscreen (Fujifilm, Japan) followed by imaging with Typhoon phosphorimager (GE Healthcare, USA). Semi-quantitative densitometric analysis of the autoradiograms was done using ImageJ software.

#### RNA extraction, reverse transcription and quantitative gene expression analysis

*Mycobacterium bovis* BCG cultures harboring either pTic6 vector alone or containing the *narL* gene were grown in 10 ml media till the mid-log phase. Expression of NarL was induced with 50 ng/mL anhydrotetracyline (aTC) for 8 hours. Total RNA was extracted using the RNeasy Mini Kit (Qiagen, USA) according to the manufacturer’s prescribed protocol from the exponentially grown cultures. RNA yield was quantified using the NanoDrop® ND-1000 UV-Vis Spectrophotometer (NanoDrop Technologies, USA). 1μg of purified RNA was used to make cDNA using the iScript™ cDNA synthesis kit (Bio-Rad, USA) using the manufacturer’s protocol. Quantitative real time PCR (qRT-PCR) was performed with 1μl of the cDNA reaction using the SYBR Flash in the Rotor-Gene Q (Qiagen, USA) according to the manufacturer’s protocol.

## Supporting information

Supplementary Material for Sequestration of histidine kinase by non-cognate response regulators establishes a threshold level of stimulation for TCS

## ACKNOWLEDGEMENTS

This work was supported by the Wellcome Trust/DBT India Alliance Senior Fellowship IA/S/14/1/501307 (NMD). Department of Biotechnology, India (Grant No. BT/PR17357/MED/29/1019/2016) to DKS. The study is also supported in part by the DBT partnership program to Indian Institute of Science (DBT/PR27952-INF/22/212/2018). Equipment support by DST– Funds for Infrastructure in Science and Technology program (SR/FST/LSII-036/2016). We thank Prof. Hemalatha Balaram (JNCASR) for help with MST, Dr. Sivaramaiah Nallapeta and Saji Menon (Nanotemper, India) for support with the MST instrument and the MO Affinity Analysis software.

## AUTHOR CONTRIBUTIONS

GDS performed the experiments and analysis; RR did the mathematical modeling and analysis; GDS and RR wrote the first drafts; DKS and NMD contributed to the analysis and revisions of the manuscript.

