## Supplementary Material for Sequestration of histidine kinase by non-cognate response regulators establishes a threshold level of stimulation for TCS for "Sequestration of histidine kinases by non-cognate response regulators establishes a threshold level of stimulation for bacterial two-component signaling"

**TABLE S1. Model parameters and their estimates**

| Symbol | Meaning | Value (95% CI) | Units | Source |
|---|---|---|---|---|
| $k_d^{lig}$ | Ligand binding rate constant | $6\times10^4$ | min$^{-1}$ | Ref. 1 |
| $k_f^{lig}$ | Ligand dissociation rate constant | $6\times10^3$ | min$^{-1}$ | Ref. 1 |
| $k_f^{ATP}$ | ATP binding rate constant | 0.06 | nM$^{-1}$min$^{-1}$ | Ref. 2 |
| $K_E$ | ATP equilibrium association constant | $1.7\times10^{-6}$ | nM$^{-1}$ | Ref. 2 |
| $k_p$ | Autophosphorylation rate constant | 0.27 | min$^{-1}$ | Ref. 2 |
| $k_f^{RR}$, $k_f^{RR-nc}$ | Binding rate constant of HK with cognate and non-cognate RR | 0.06 | nM$^{-1}$min$^{-1}$ | Ref. 1 |
| $K_D$ | Equilibrium dissociation constant of HK-$RR_c$ binding | 268 | nM | Present study (Fig. 1) |
| $k_f^{tc}$ | Rate constant of transition complex formation | 100 | min$^{-1}$ | Assumed based on fast dynamics |
| $K_{tc}$ | Equilibrium dissociation constant of transition complex | $1.99\ (1.31\ -\ 3.02)$ | – | Present study (Fig. 2) |
| $k_p^{RR}$ | Phosphotransfer rate constant | $1.83\ (1.04\ -\ 3.33)$ | min$^{-1}$ | Present study (Fig. 2) |
| $k_{dp}^{RR}$ | Binding rate constant of HK and $RR_c$ | $8\times10^{-4}\ (5\times10^{-4}\ -\ 14\times10^{-4})$ | nM$^{-1}$min$^{-1}$ | Present study (Fig. 2) |

| | | | | |
|---|---|---|---|---|
| $k_d^{tc}$ | Rate constant of dissociation of transition complex | $0.57\ (0.32\ -\ 1.00)$ | $min^{-1}$ | Present study (Fig. 2) |
| $K_{D-nc}$ | Equilibrium dissociation constant of HK-$RR_{nc}$ | 80 | nM | Present study (Fig. 1) |
| $k_d^{tc-nc}$ | Rate constant of dissociation of transition complex containing $RR_{nc}$ | $0.10\ (0.05\ -\ 0.20)$ | $min^{-1}$ | Present study (Fig. 2) |
| $k_d^{RR}$ | Dephosphorylation rate constant of $RR_c^*$ | 0.0144 | $min^{-1}$ | Ref. 3 |
| $P_T$ | Total promotor binding regions | 100 | nM | Ref. 1 |
| $\alpha$ | Fold-increase in transcription due to $RR_c^*$ binding | 10 | – | Ref. 4 |
| $\beta$ | Effective rate constant of $RR_c$ synthesis | 0.36 | $nM\,min^{-1}$ | Ref. 4 |
| $\lambda$ | Rate of HK synthesis relative to $RR_c$ synthesis | 0.1 | – | Ref. 4 |
| $K$ | Equilibrium association constant of $RR_c$ with promoter | $5\times10^5$ | $nM^2$ | Ref. 4 |
| $k_{\deg}$ | Degradation rate constant of proteins | $4.17\times10^{-4}$ | $min^{-1}$ | Ref. 5 |
| $\phi_{HK}$ | Fraction of reaction competent HK | 0.14 | – | Ref. 2 |

| | | | | |
|---|---|---|---|---|
| $\phi_{RR}$ | Fraction of reaction competent $RR_c$ | $0.036\ (0.028\ -\ 0.045)$ | – | Present study (Fig. 2) |

### SUPPLEMENTARY FIGURES

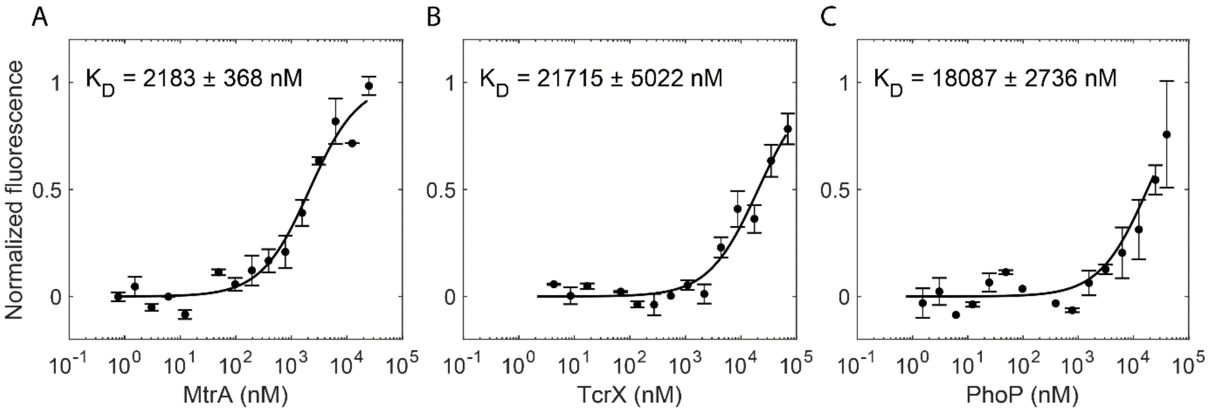


**Figure S1. Binding affinities of unphosphorylated MtrB for cognate and non-cognate RRs.** Changes in the thermophoretic movement of fluorescently tagged MtrB were measured as a function of titrant concentration as described in the methods section. Interaction analysis of 50 nM of MtrB-GFP with titrant RRs (range): **(A)** MtrA (0.76 nM to 25 μM), **(B)** PhoP (1.52 nM to 40 μM), and **(C)** TcrX (4.27 nM to 70 μM). $K_D$ values were evaluated by plotting normalized fluorescence against the logarithmic concentrations of serially diluted ligand (RRs). Symbols are mean ± S.E.M. from more than 3 independent experiments and curves are best-fits.

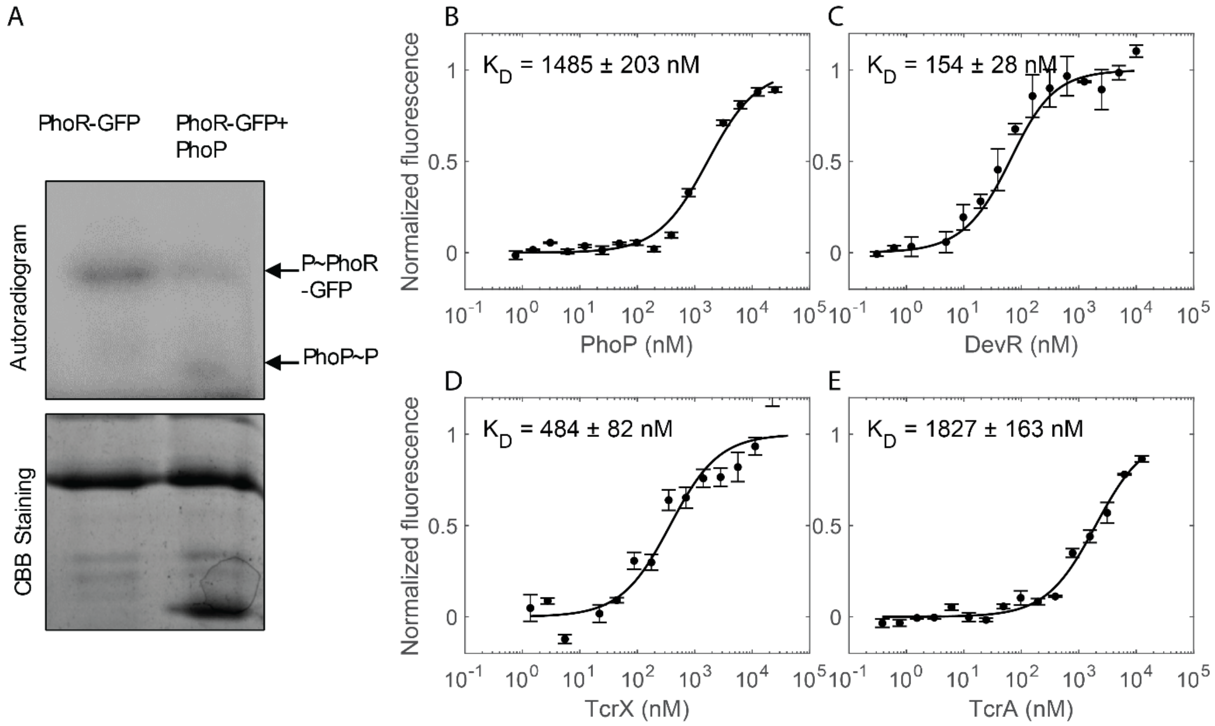


**Figure S2. Functional activity assay and binding affinities of phosphorylated PhoR-GFP for cognate and non-cognate RRs. (A)** Autophosphorylation of the phosphorylated HK PhoR-GFP with subsequent phosphotransfer to the cognate RR PhoP. Changes in the thermophoretic movement of fluorescently tagged target HK post autophosphorylation were measured as a function of titrant concentration as described in the methods section. Interaction analysis of 50 nM of P~PhoR-GFP with titrant RRs (range): (**B)** PhoP (0.73 nM to 25 μM), **(C)** DevR (0.31 nM to 10 μM), **(D)** TcrX (1.37 nM to 22.5 μM), (**E)** TcrA (0.38 nM to 12.5 μM). $K_D$ values were evaluated by plotting normalized fluorescence against the logarithmic concentrations of serially diluted ligand (RRs). Symbols are mean ± S.E.M. from more than 3 independent experiments and curves are best-fits.

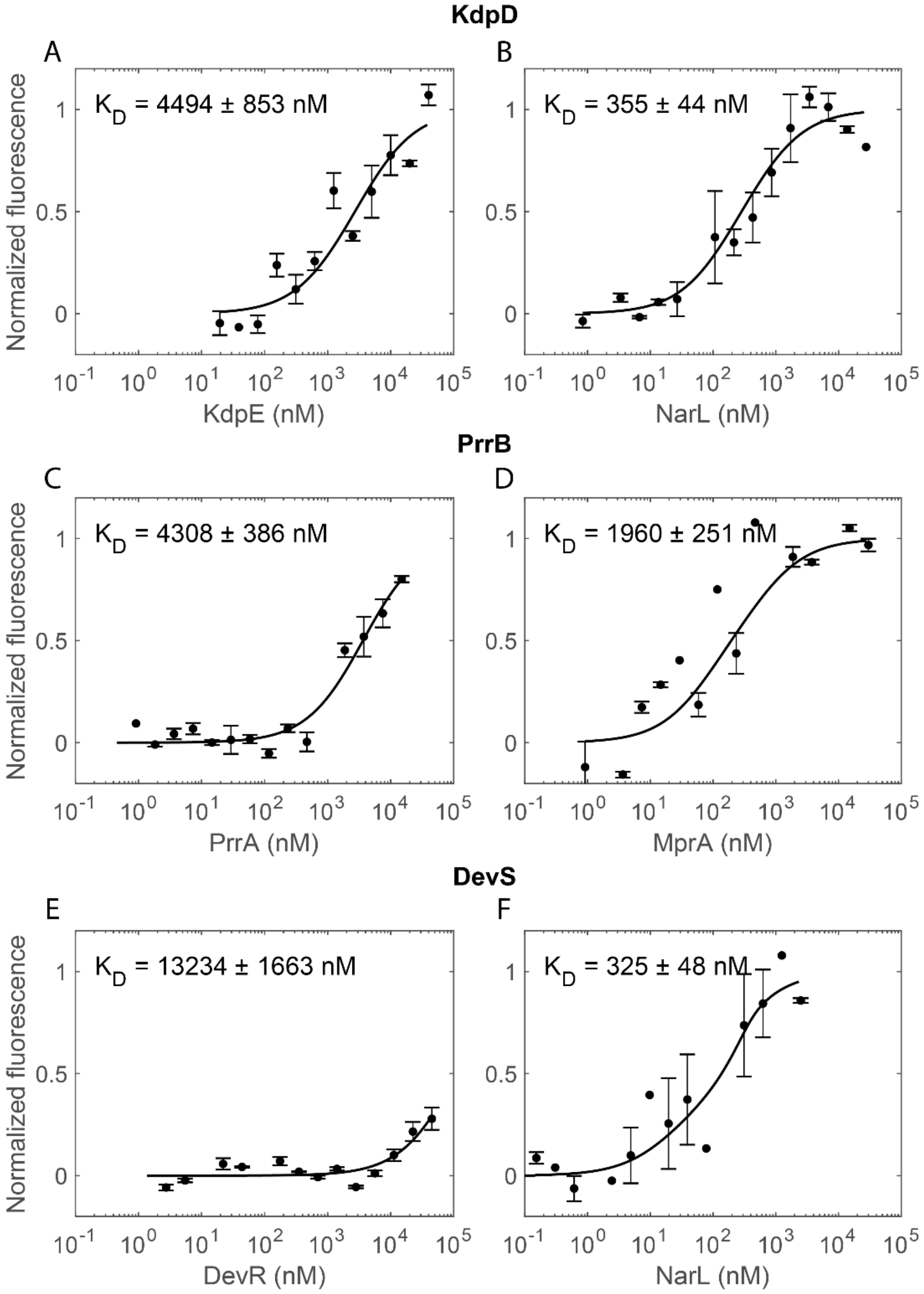


**Figure S3. Binding affinities of phosphorylated HKs for cognate and non-cognate RRs.** Interaction analysis of 50 nM of the HK P~KdpD with titrant RRs (range): **(A)** KdpE (19.5 nM to 40 µM) , **(B)** NarL (0.84 nM to 27.5 µM); of the HK P~PrrB with the titrant RRs (range) **(C)** PrrA (0.92 nM to 15 µM), **(D)** MprA (0.92 nM to 30 µM); of the HK P~DevS with the titrant RRs (range) **(E)** DevR (2.75 nM to 45 µM), and **(F)** NarL (0.15 nM to 5 µM). $K_D$ values were evaluated by plotting normalized fluorescence against the logarithmic concentrations of serially diluted ligand (RRs). Symbols are mean ± S.E.M. from more than 3 independent experiments and curves are best-fits.

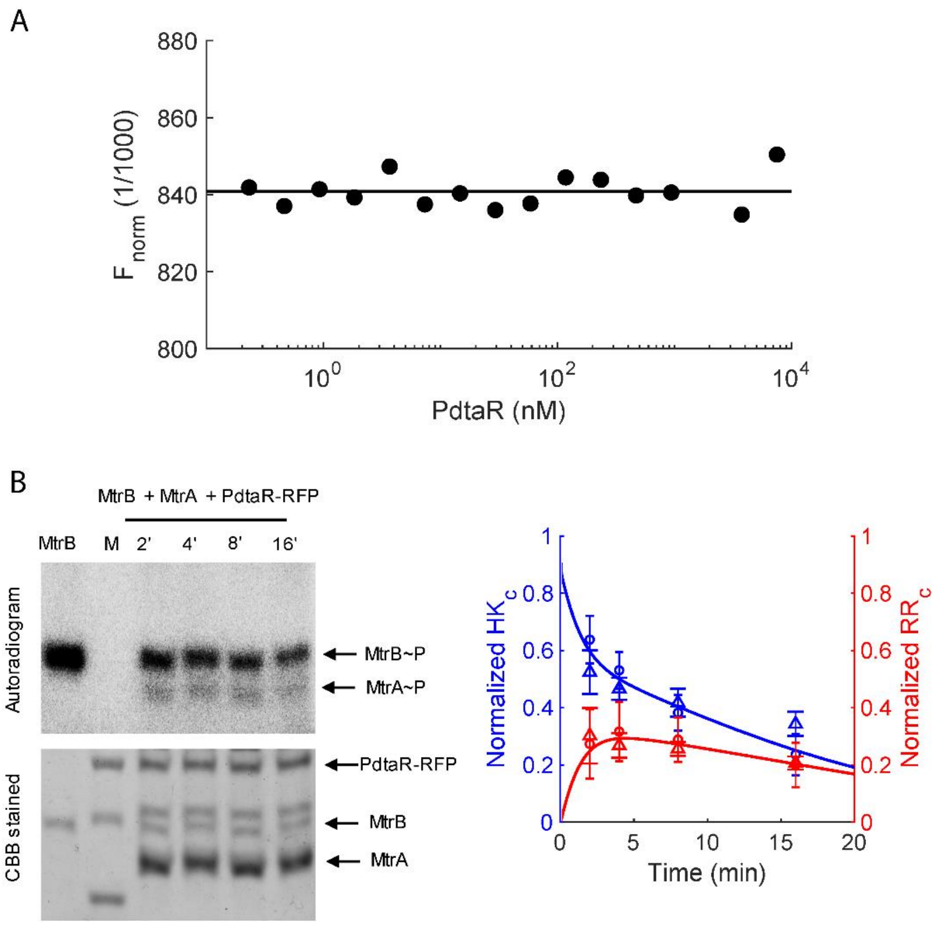


**Figure S4. Phosphotransfer kinetics from MtrB to MtrA with non-binding PdtaR-RFP. (A)** Representative thermophoretic change profile of 50 nM of P~MtrB-GFP with the titrant RR PdtaR (0.23 nM to 7.5 μM), indicating no binding. **(B)** Time course assay of the phosphotransfer from the HK MtrB~P to the cognate RR MtrA in the presence of the non-cognate RR PdtaR-RFP followed by densitometric analysis of the time course assays on the right side. The autophosphorylation control was used to normalize the intensities of the individual bands. Blue symbols represent MtrB~P and red symbols MtrA~P. Lines represent best-fits of our model (Methods). The error bars represent mean ± S.E.M (n=3).

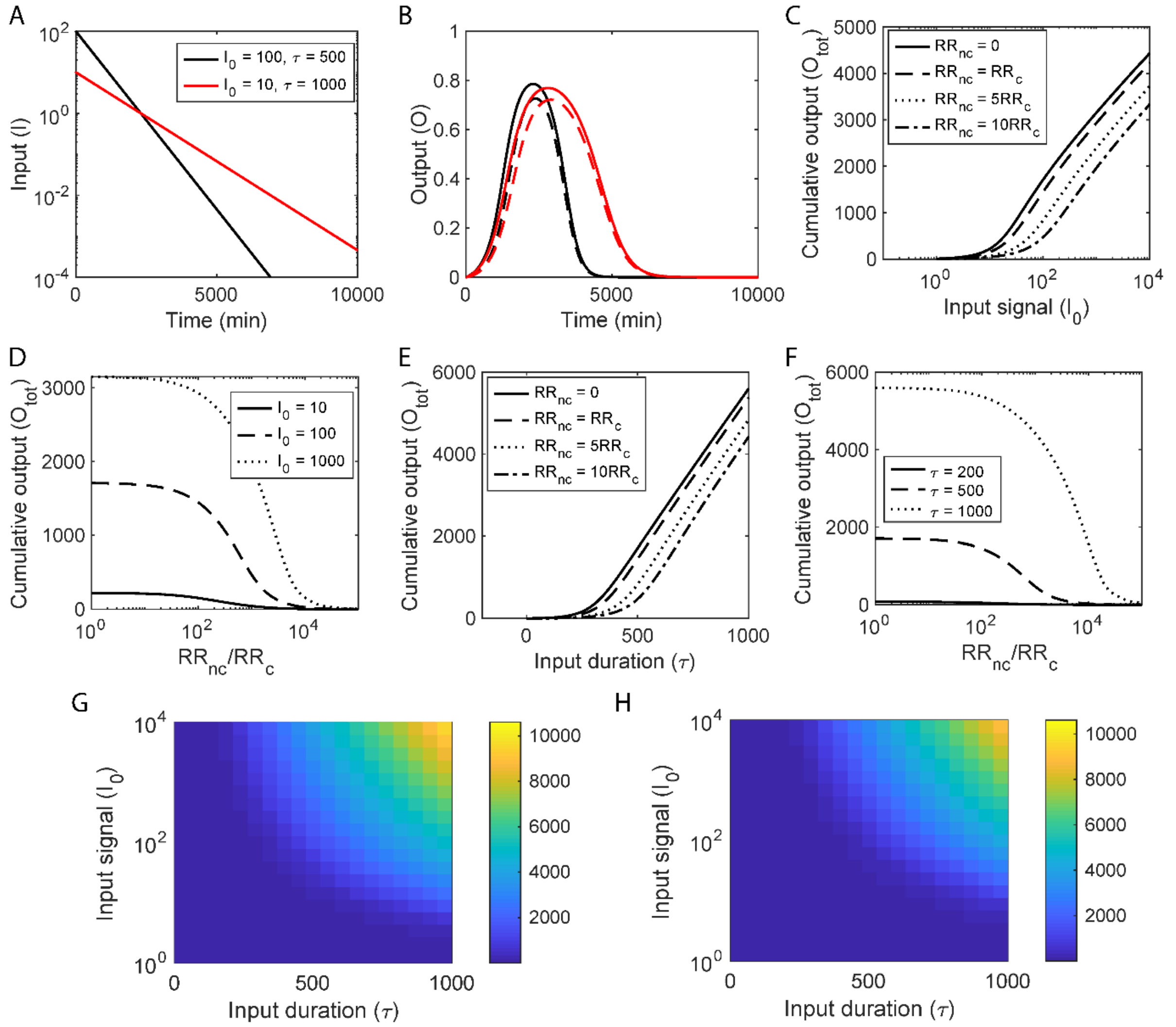


**Figure S5. Model predictions of the total TCS signal response ($O_{total}$) and the impact of sequestration. (A)** Representative inputs, $I$, indicating strong but short-lived (black) and weak but extended (red) stimuli. **(B)** The corresponding outputs without (solid lines) and with (dashed lines) sequestration by a non-cognate RR. **(C)** The total response ($O_{total}$) as a function of the maximum input, $I_0$, for different extents of sequestration, determined by the ratios of the non-cognate RRs, $RR_{nc}$ to the cognate RR, $RR_c$, indicated. **(D)** $O_{total}$ as a function of the ratio $RR_{nc}/RR_c$ for different $I_0$. **(E)** $O_{total}$ as a function of the signal half-life, $\tau$, for different values of $RR_{nc}/RR_c$. **(F)** $O_{total}$ as a function of $RR_{nc}/RR_c$ for different values of $\tau$. ($\tau$ is in minutes throughout.) Heatmaps showing $O_{total}$ as functions of $I_0$ and $\tau$ in the **(G)** absence or **(H)** presence of non-cognate RRs, indicating the threshold stimulation for response shifting to higher $I_0$ and $\tau$ with sequestration. Corresponding calculations for the peak response, $O_{max}$, are in Figure 5. Model predictions were obtained by solving Eqs. (1)-(33) (Methods) using parameter values listed in Table S1.
